# The zebrafish orthologue of familial Alzheimer’s disease gene *PRESENILIN* 2 is required for normal adult melanotic skin pigmentation

**DOI:** 10.1101/414144

**Authors:** Haowei Jiang, Morgan Newman, Michael Lardelli

**Author notes:** **Corresponding Author:** Haowei Jiang, University of Adelaide, School of Biological Sciences, Alzheimer’s Disease Genetics Laboratory, North Terrace, Adelaide, SA 5005, AUSTRALIA.

## Abstract

Alzheimer’s disease is the most common form of age-related dementia. At least 15 mutations in the human gene *PRESENILIN 2* (*PSEN2*) have been found to cause familial Alzheimer’s disease (fAD). Zebrafish possess an orthologous gene, *psen2*, and present opportunities for investigation of *PRESENILIN* function related to Alzheimer’s disease. The most prevalent and best characterized fAD mutation in *PSEN2* is *N141I*. The equivalent codon in zebrafish *psen2* is N140. We used genome editing technology in zebrafish to target generation of mutations to the N140 codon. We isolated two mutations: *psen2*^*N140fs*^, (hereafter “*N140fs*”), causing truncation of the coding sequence, and *psen2*^*T141_L142delinsMISLISV*^, (hereafter “*T141_L142delinsMISLISV*”), that deletes the two codons immediately downstream of N140 and replaces them with seven codons coding for amino acid residues MISLISV. Thus, like almost every fAD mutation in the *PRESENILIN* genes, this latter mutation does not truncate the gene’s open reading frame. Both mutations are homozygous viable although *N140fs* transcripts are subject to nonsense-mediated decay and lack any possibility of coding for an active γ-secretase enzyme. *N140fs* homozygous larvae initially show grossly normal melanotic skin pigmentation but subsequently lose this as they grow while retaining pigmentation in the retinal pigmented epithelium. *T141_L142delinsMISLISV* homozygotes retain faint skin melanotic pigmentation as adults, most likely indicating that the protein encoded by this allele retains weak γ-secretase activity. Null mutations in the human *PRESENILIN* genes do not cause Alzheimer’s disease so these two mutations may be useful for future investigation of the differential effects of null and fAD-like *PRESENILIN* mutations on brain aging.

**Financial Disclosure Statement:** This research was supported by grants from the National Health and Medical Research Council of Australia, GNT1061006 and GNT1126422, and by funds from the School of Biological Sciences of the University of Adelaide. HJ is supported by an Adelaide Scholarship International from the University of Adelaide.

**Conflict of Interest Statement:** The authors declare no conflict of interest.

## Introduction

Alzheimer’s disease (AD) is a progressive neurodegenerative disorder, and is the most common form of age-related dementia, accounting for 50-75% of dementia cases worldwide (1). Most AD occurs after the age of 65 years (late onset) and is sporadic. Early onset AD is far less common and approximately 13% of early onset cases are familial AD (fAD) (2). Autosomal dominant inheritance of mutations in the *AMYLOID BETA A4 PRECURSOR PROTEIN* gene (*APP*) (3), *PRESENILIN 1* and *2* genes (*PSEN1, PSEN2*) (4), and *SORTILIN-RELATED RECEPTOR* gene (*SORL1*) (5, 6) are considered to be the major cause of fAD. Of the two *PRESENILIN* genes, *PSEN2* is a less common locus for fAD mutations than *PSEN1*. Only around 15 fAD mutations have been reported in *PSEN2* to date, compared to over two hundred mutations reported in *PSEN1* (4). All but one of the many different fAD mutations in the *PSEN* genes do not cause truncation of coding sequences, a phenomenon we have previously described as the “fAD mutation reading frame preservation rule” (7).

PSEN proteins become endoproteolytically cleaved during activation of γ-secretase activity to form N- and C-terminal fragments (NTF and CTF resprectively) (8). The NTFs and CTFs of PSEN2 predominantly localise to the endoplasmic reticulum (ER) (9) and to late endosomes / lysosomes (10). The first two transmembrane domains (TMDs) of PSEN2 are thought to be necessary for ER localisation (11) while a conserved sequence near the N-terminal is bound by Adaptor Complex AP-1 to direct PSEN2 protein to late endosomes / lysosomes (10). The localisation of PSEN2, rather than PSEN1, to late endosomes / lysosomes implies a particular importance for PSEN2 in the biogenesis of melanosomes (10, 12), an organelle type related to lysosomes (13) that is specialised for formation of the dark pigment melanin (14).

The first fAD mutation reported in *PSEN2* was *N141I*, caused by an A-to-T transition at the second position of codon 141 (15). The *N141I* mutation alters the N-terminal flank of the second TMD (TMD2) of PSEN2 by substituting a hydrophobic isoleucine residue for the hydrophilic asparagine residue immediately downstream of the first residue of TMD2. This position is thought to be important for accurate positioning of the transmembrane α-helix structure (16). A PolyPhen-2 (17) analysis of the *N141I* mutation indicates probable damage to protein structure with a score of 0.934 (sensitivity: 0.80; specificity: 0.94). The mean age of Alzheimer’s disease onset for carriers of *N141I* is 53.7 years old, but with a very wide range of 39 to 75 years (4). Thus, *N141I* has an age of onset overlapping those of *PSEN1* fAD families (mean age of onset of 45.5 years) and sporadic AD (mean age of onset of 71.5) (4). The *N141I* mutation is thought to increase the ratio of Aβ42 to Aβ40 via abnormal γ-secretase activity (18). A more recent transgenic mouse model of AD suggested that both Aβ42 and Aβ40 production are enhanced by *N141I*, and this can signficantly accelerate Aβ-dependent dysfunction in spatial learning and memory (19).

Mammalian PRESENILINs have also been found necessary for tyrosinase trafficking and melanin formation by a γ-secretase-dependent mechanism (20). TYROSINASE is a key enzyme in melanin synthesis (21). The two TYROSINASE-related proteins, TYROSINASE-related protein 1 (Tyrp1) and DOPACHROME TAUTOMERASE (DCT) (also known as TYROSINASE-related protein 2 (Tyrp2)) (22), are implicated in the activity of the intramembrane protease, γ-secretase (20, 23). A partial loss-of-function in melanotic pigment formation has been observed in a mouse model of the *PSEN1* fAD mutation *M146V* (20).

In mammals, the protein SILVER, MOUSE, HOMOLOG OF (SILV, also known as PREMELANOSOMAL PROTEIN, PMEL) (24) is another type 1 membrane protein that can be cleaved by proteases including γ-secretase (25) to form a natural functional amyloid that facilitates melanin formation (12). SILV is expressed in pigment cells of the eye and skin, which synthesise melanin pigments within melanosomes (26). After a juxtamembrane cleavage, the C-terminal fragment of SILV is then processed by the γ-secretase complex to release an intracellular domain fragment (25) into endosomal precursors to form amyloid fibrils. These ultimately become melanosomes (27, 28).

Zebrafish are a versatile system in which to investigate, at the molecular level, the effects on the brain and other tissues of fAD mutations (29). The ability to generate large families of siblings and then raise these in a near identical environment (the same tank or the same recirculated-water system) can reduce genetic and environmental variability to allow more sensitive detection of mutation-dependent changes. The organisation of the genome and the genetic pathways controlling signal transduction and development of zebrafish and humans are highly conserved (30). Despite ∼420 million years of divergent evolution of the human and zebrafish lineages (31), most human genes have clearly identifiable orthologues in zebrafish. Thus, the zebrafish genes *psen1* (32) and *psen2* (33) are orthologues of human *PSEN1* and *PSEN2*, respectively. The Presenilin protein sequences of zebrafish show considerable identity with those of humans. The zebrafish Psen1 protein shows 73.9% amino acid residue (aa) identity with human PSEN1 (32), while zebrafish Psen2 shows 74% identity with human PSEN2 (33).

In this paper, we describe an attempt to generate a zebrafish model of the *N141I* fAD mutation of human *PSEN2* by introducing an equivalent mutation into the zebrafish *psen2* gene. While homology-directed repair (HDR) after CRISPR Cas9 cleavage at the relevant site in zebrafish *psen2* was not successful, we did find products of non-homologous end joining (NHEJ) that will prove useful in future analyses. We identified both a frameshift mutation and a reading frame-preserving indel mutation close to the N141-equivalent codon of zebrafish *psen2* (N140). Surprisingly, we discovered that the γ-secretase activity of Psen2 (unlike that of Psen1) appears essential for melanotic pigment formation in the skin of zebrafish adults but not in their retinal pigmented epithelium.

## Materials and Methods

### Animal ethics

All experiments using zebrafish were conducted under the auspices of the Animal Ethics Committee of the University of Adelaide. Permits S-2014-108 and S-2017-073.

### CRISPR guide RNA (sgRNA) design and synthesis

The target sequence of the sgRNA used to generate double-stranded breaks near the N140 codon in zebrafish *psen2* is 5’-GAATTCGGTGCTCAACACTC *TGG*-3’. The template for sgRNA transcription was synthesised by PCR (34). The forward primer for this template synthesis PCR contains a T7 polymerase binding site (the underlined region), the target sequence (bold) and a region complementary to a common reverse primer (italicised): 5’-<u>GAAATTAATACGACTCACTATAGG</u>**GAATTCGGTGCTCAACACTC***GTTTT AGAGCTAGAAATAGC*-3’. The sequence of the reverse primer is 5’- AAAAGCACCGACTCGGTGCCACTTTTTCAAGTTGATAACGGACTAGCCTTA TTTTAACTTGCTATTTCTAGCTCTAAAAC-3’. This synthesis PCR used Phusion^®^ High-Fidelity DNA Polymerase (NEB, Ipswich, Massachusetts, USA, M0530S) and cycle conditions of 98°C for 30 s and then 35 cycles of [98°C, 10 s; 60°C, 30 s; 72°C, 15 s] then 72°C, 10 min. The template was then gel-purified using the Wizard^®^ SV Gel and PCR Clean-Up System (Promega, Madison, Wisconsin, USA, A9281). The target sgRNA was synthesized from this template using the HiScribe™ T7 Quick High Yield RNA Synthesis Kit (NEB, Ipswich, Massachusetts, USA, E2050S).

### Design of single-stranded oligonucleotide templates for homology-directed repair (HDR)

To attempt to introduce the *N140I* mutation into zebrafish *psen2* (equivalent to human *PSEN2 N141I*), a single stranded oligonucleotide template (“N140I oligo”) containing the N>I mutation (A>T, bold italics and underlined) followed by two silent (synonymous codon) mutations (T>C and G>C, italicised and underlined) was designed: 5’-ACTCAGTGGGCCAGCGTCTGCTGAATTCGGTGCTC***A<u>T</u>C****AC<u>C</u>CT<u>C</u>*GTCATG ATCAGTGTGATTGTCTTCATGACC-3’.

We also attempted (unsuccessfully) to introduce the *V147I* mutation into zebrafish *psen2*, (equivalent to *V148I* in human *PSEN2*) using a single-stranded oligonucleotide template (“V147I oligo”), containing the V>I mutation (G>A and G>C, bold italics and underlind) followed by two silent (synonymous codon) mutations (T>A and C>G, italicised and underlined): 5’- CTGAATTCGGTGCTCAACACTCTGGTCATGATCAGTA***T***C*AT<u>A</u>GT<u>G</u>*TTCATGA CCATCATCCTGGTGCTGCTCTAC-3’. The attempted mutation of the *V147* site in *psen2* is only described and discussed in Supplemental Information.

The single-stranded oligonucleotide templates were co-injected with their corresponding CRISPR/Cas9 systems, so that any induced double-stranded DNA breaks (DSBs) might be repaired through the HDR pathway (35) to insert desired mutations into the zebrafish genome.

### Injection of zebrafish embryos

Tübingen (TU) strain wild type embryos were collected from mass spawning. The target sgRNA (70 ng/μL final concentration) was mixed with “N140I oligo” (30 ng/μL for final concentration) and Cas9 nuclease (1μg/μL for final concentration) (Invitrogen, Carlsbad, California, USA, B25640), and then incubated at 37°C for 15 min to maximize formation of active CRISPR Cas9 complexes. 5-10 nL of the mixture was then injected into zebrafish embryos at the one-cell stage. The injected embryos were subsequently raised for mutation screening.

### Mutation detection in CRISPR Cas9-injected G0 fish

From each batch of injected embryos, 10 embryos were selected at random at ∼24 hpf and pooled for genomic DNA extraction. The genomic DNA of these embryos was extracted using sodium hydroxide (36). The 10 embryos were placed in 100 μL of 50 mM NaOH and then heated to 95°C for 15 min. They were then cooled to 4°C followed by addition of 1/10th volume of 1 M Tris-HCl, pH 8.0 to neutralize the basic solution (36).

Mutation-specific primers were designed to detect mutation-carrying fish by PCR. For the “N140I oligo”-injected embryos, a mutation-specific forward primer was designed: 5’-TCGGTGCTC***A<u>T</u>C****AC<u>C</u>CT<u>C</u>*-3’. A wild type-specific forward primer (5’-TCGGTGCTC*AACACTCTG*-3’) and a common reverse primer (5’-ACCAAGGACCACTGATTCAGC-3’) were also designed. The PCR conditions for both these reactions are: 95°C, 2 min and then 31 cycles of [95°C, 30 s; 58°C, 30 s; 72°C 30 s], then 72°C, 5 min. The lengths of the expected PCR products of these reactions are all ∼300 nucleotides.

For the “V147I oligo”-injected embryos, a mutation-specific forward primer was designed: 5’-TCTGGTCATGATCAGT***<u>A</u>T<u>C</u>****AT<u>A</u>GT<u>G</u>*-3’. A wild type-specific forward primer (5’-TCTGGTCATGATCAGT*GTGATTGTC*-3’) and a common reverse primer (5’-TCACCAAGGACCACTGATTCAGC-3’) were also designed. The PCR conditions for all these three reactions are: 95°C, 2 min, and then 31 cycles of [95°C, 30 s; 58°C, 30 s; 72°C, 30 s], then 72°C 5 min. The lengths of the PCR products of these reactions are ∼280 nucleotides.

The F1 progeny of the mosaic, mutation-carrying G0 fish were also screened with these mutation-specific PCR reactions.

### Mutation detection in F1 fish using the T7 endonuclease I assay

Since the DSBs induced by the CRISPR/Cas9 system may also be repaired through the NHEJ pathway (35, 37), random mutations may also be generated at the DSB sites. Thus, the F1 progeny of the mosaic, mutation-carrying G0 fish may be heterozygous for such random mutations.

To screen for these mutations, the genomic DNA of tail biopsies from F1 fish was extracted using sodium hydroxide as above, followed by analysis using the T7 endonuclease I assay (since T7 endonuclease I is able to recognize and cleave at the sites of mismatches in DNA heteroduplexes (38)).

A pair of amplification primers binding in the regions flanking the N140 target site was designed: 5’-AGCATCACCTTGATTCAAGG-3’ and 5’-GGTTCCTGATGACACACTGA-3’. The PCR conditions for this amplification reaction are 95°C, 2 min and then 31 cycles of [95°C, 30 s; 58°C, 30 s; 72°C, 30 s], then 72°C, 5 min and the amplified fragment is predicted to be 473 nucleotides in length. The PCR products were purified using the Wizard^®^ SV Gel and PCR Clean-Up System (Promega, Wisconsin, USA, A9281). These purified PCR products were then denatured at 95°C for 5 min and then annealed by slow cooling of the samples at −2°C/sec from 95°C to 85°C and then −0.1°C/sec from 85°C to 25°C). Finally, the annealed PCR products were digested using T7 endonuclease I (NEB, Ipswich, Massachusetts, USA, M0302S) was added to. If reannealed fragments contained mismatches due to mutations, they would be cleaved by T7 endonuclease I into two fragments; ∼109 nucleotides (upstream) and ∼364 nucleotides (downstream). Those amplified and reannealed fragments showing positive signals (cleavage) in T7 endonuclease I assays were then sent to the Australian Genome Research Facility (AGRF, North Melbourne, VIC, Australia) for **S**anger sequencing to identify the mutations.

### Mutation detection in F2 fish using PCR

Mutation-specific PCR primers were designed to detect the two mutations (*N140fs* and *T141_L142delinsMISLISV*) identified in F1 fish. For *N140fs*, a mutation-specific forward primer (5’-TGCTGAATTCGGTGCTCTG-3’) was designed. For *T141_L142delinsMISLISV*, another mutation-specific forward primer (5’- TGAATTCGGTGCTCAACATG-3’) was designed. A wild type-specific forward primer (5’-TGAATTCGGTGCTCAACACTC-3’) was designed as a control. A common reverse primer (5’-TCACCAAGGACCACTGATTCAGC-3’) was used with these three different forward primers. The temperature cycling conditions for these PCRs are identical for the wild type and *N140fs* alleles: 95°C, 2 min, and then 31 cycles of [95°C, 30 s; 60°C, 30 s; 72°C, 30 s], then 72°C, 5 min. For detection of the *T141_L142delinsMISLISV* allele, the annealing temperature was altered to 61.5°C. The PCR products of these reactions are all predicted to be ∼320 nucleotides in length.

### Breeding of mutant fish

Since the mutation-carrying G0 fish were mosaic for mutations, these were outbred with wild type TU fish so that their progeny (F1 fish) would be completely heterozygous for any mutations.

The F1 fish carrying the *T141_L142delinsMISLISV* or *N140fs* alleles were outbred with wild type TU fish to generate additional individuals heterozygous for the mutations. (The families of progeny of such matings would consist of 50% heterozygous mutants and 50% wild type fish). When these F2 progeny were sexually mature, pairs of heterozygous individuals were in-crossed to generate F3 families containing homozygous mutant, heterozygous mutant and wild type siblings for further analysis.

### Imaging of skin pigmentation in zebrafish

The pigmentation patterns of mutant zebrafish were imaged using a Leica Microsystems, Type DFC450 C microscope, and the software Leica Application Suite, Version 4.9.0 (Leica Microsystems, Wetzlar, Germany).

### Total RNA extraction from 6-month-old zebrafish brains

When F2 fish families from outcrossed heterozygous mutant F1 fish were 6 months of age, eight female fish of each genotype (i.e. eight wild type and eight heterozygous mutant individuals) were selected for brain removal (after euthanized by submersion in ice water) and total RNA extraction. From these fish, four of each genotype were exposed to hypoxia according to the method we have previously established (39). The dissolved oxygen content of the hypoxic water was ∼1.00 mg/L (treated for ∼2.5 h), while the other four fish of each genotype were exposed to normoxia (i.e. the dissolved oxygen content of the normoxic water was ∼6.60 mg/L). Total RNA was extracted from these brains using the *mir*Vana™ miRNA Isolation Kit (Ambion, Inc, Foster City, California, USA, AM1560). cDNA was synthesised from the RNA using the SuperScript™ III First-Strand Synthesis System (Invitrogen, Carlsbad, California, USA, 18080051) and Random Primers (Promega, Madison, Wisconsin, USA, C1181).

### Allele-specific expression analysis by digital quantitative PCR (dqPCR)

PCR primer pairs detecting specific alleles were designed for dqPCR: a specific forward primer for mutation *T141_L142delinsMISLISV* (5’-TGAATTCGGTGCTCAACATG-3’), a specific forward primer for mutation *N140fs* (5’-TGCTGAATTCGGTGCTCTG-3’), and a specific forward primer for the wild type allele (5’-TGAATTCGGTGCTCAACACTC-3’). A common reverse primer (5’-AAGAGCAGCATCAGCGAGG-3’) was used with all these three forward primers. Allele-specific dqPCR was performed using the QuantStudio™ 3D Digital PCR System (Life Sciences, Waltham, MA, USA) with QuantStudio™ 3D Digital PCR 20K Chip Kit v2 and Master Mix (Life Sciences, Waltham, MA, USA, A26317) and SYBR™ Green I Nucleic Acid Gel Stain (Life Sciences, Waltham, MA, USA, S7563). The dqPCR conditions for allele-specific expression detection are 96°C, 10 min, then 49 cycles of [62°C, 2 min; 98°C, 30 s], then 62°C 2 min. The expected length of the PCR products is ∼130 bp. 25ng of cDNA (based on quantification of RNA concentration and the assumption of complete reverse transcription into cDNA) of each sample was loaded into each chip. The chips were analysed using QuantStudio™ 3D AnalysisSuite Cloud Software (Life Sciences, Waltham, MA, USA).

## Results

### fAD-like and coding sequence-truncating mutations in *psen2*

Our initial aim was to create mutations in zebrafish *psen2* equivalent to the fAD mutations of human *PSEN2, N141I* and *V148I* (*N140I* and *V147I* in zebrafish respectively). However, while CRISPR Cas9-targetting of these sites appeared feasible, no incorporation of desired mutations via homology-directed repair was found. Nevertheless, two mutations at the N140 site were ultimately identified. One of these is an indel mutation removing two codons (T141 and L142) and replacing these with seven novel codons (MISLISV). Consequently, this allele is designated *T141_L142delinsMISLISV* and may be considered EOfAD-like in that it does not truncate the coding sequence (CDS) (Fig 1). The second mutation is a deletion of 7 nucleotides causing a frameshift that does truncate the coding sequence, *N140fs*, due to a premature termination codon (PTC) at the 142^nd^ codon position.

**Fig 1.**
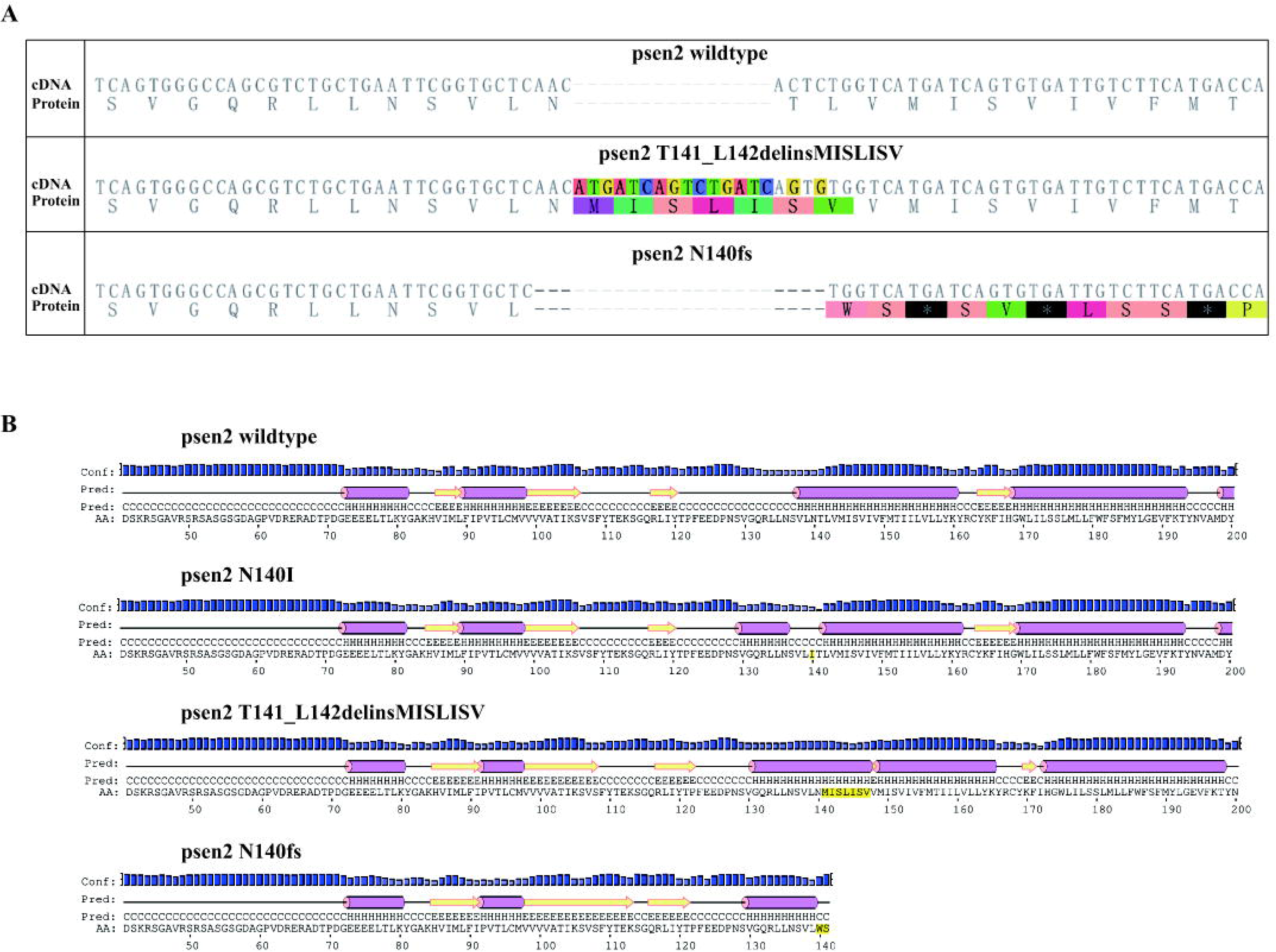
Predicted protein primary and secondary structures. (A) The protein coding sequence of zebrafish Psen2 is altered by the mutations. (B) The predicted protein structures of zebrafish Psen2 are also changed by the two identified mutations (and are shown relative to the wild type structure and a structure incorporating a hypothetical *N140I* mutation. Purple bar: helix; yellow arrow: strand; black line: coil; Conf: confidence of prediction; Pred: predicted secondary structure; AA: target sequence.

Inbreeding of *T141_L142MISLISV* and *N140fs* mutant fish showed both mutations to be homozygous viable although both showed severe defects in skin pigmentation in post-larval stages (described later).

### Changes of protein structure caused by the mutations

PRESENILINs have a complex structure with multiple TMDs. Therefore, mutations have the potential to greatly disturb protein structure by interfering with normal membrane insertion. To understand the possible consequences of, in particular, the *T141_L142delinsMISLISV* mutation, we compared theoretical hydropathicity plots (40) for our isolated mutations with those for wild type *psen2* and a mutation equivalent to human *N141I* (Fig 2). The *T141_L142delinsMISLISV* mutation contributes only non-polar (M, I, L, V) or, at least, uncharged, polar (S) amino acid residues (aa) to the protein structure, presumably expanding the hydrophobic stretch of aas that form TMD2. Presumably, this mutation allows overall correct membrane insertion but disrupts the conformation of the protein sufficiently to almost entirely, but not completely, destroy its γ-secretase activity (see later).

**Fig 2.**
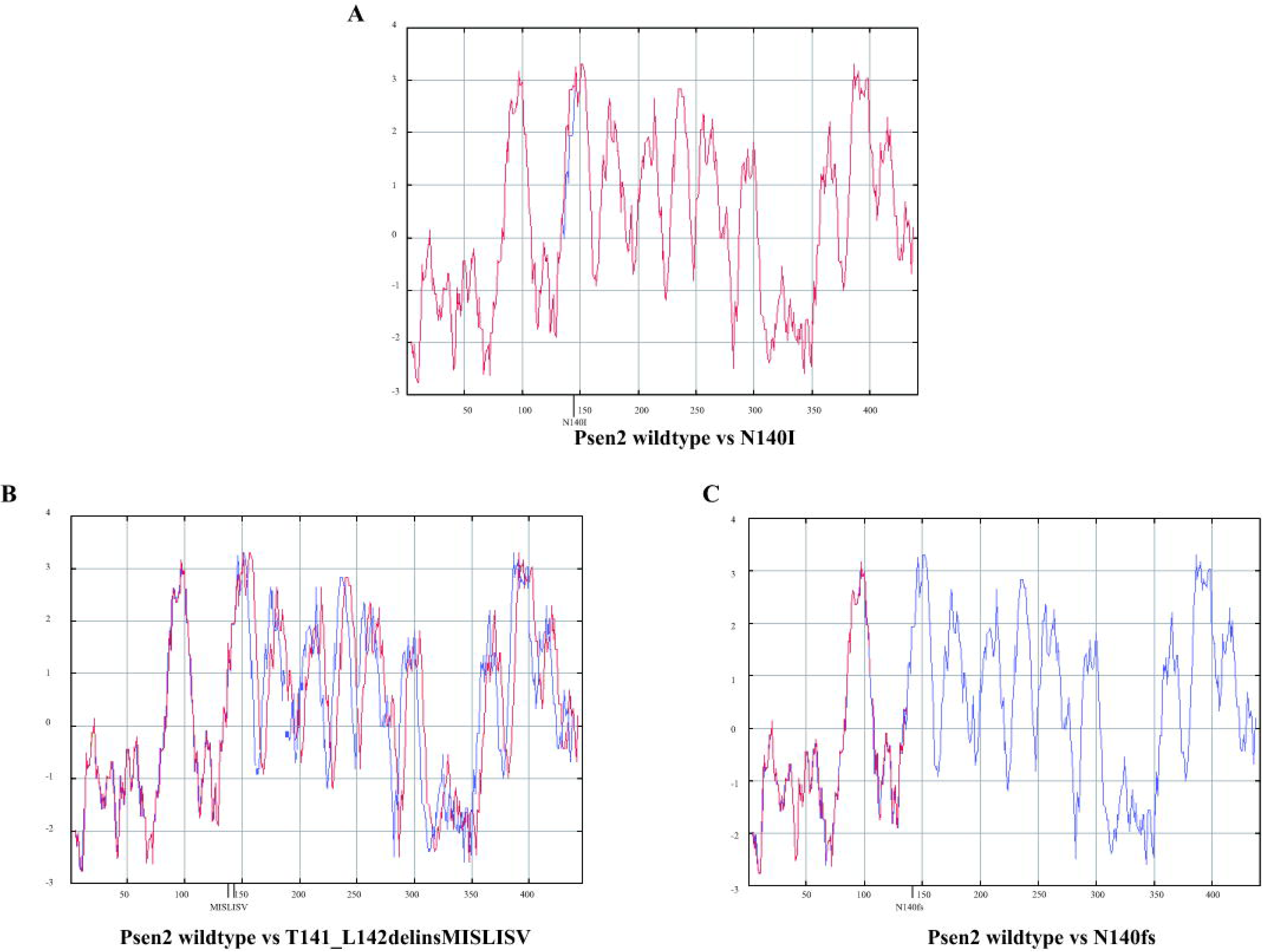
Predicted protein hydropathicity plots. The blue line refers to wild type Psen2. The red lines refer to the mutants.

The *N140fs* mutation cannot possibly express a catalytically active γ-secretase enzyme since it truncates the CDS at the start of TMD2. Thus, it lacks both the aspartate residues required for the γ-secretase catalytic domain (41, 42).

### *N140fs* transcripts are subject to nonsense-mediated decay

Mutations creating premature termination codons (PTCs) in coding sequences upstream of exon-exon junctions in spliced transcripts can result in destabilisation of the transcripts through nonsense-mediated decay (NMD, reviewed by (43)). Therefore, we expected that transcripts from the *T141_L142delinsMISLISV* allele might be similarly stable to wild type transcripts while *N140fs* allele transcripts would show decreased stability and abundance. To test this we performed dqPCR that allows direct comparison of transcript abundances. We extracted total RNA from the brains of 6-month-old adult zebrafish, reverse transcribed this to cDNA, and then performed dqPCR with primers specifically detecting the wild type or mutant alleles. The results confirmed similar levels of *T141_L142delinsMISLISV* and wild type transcripts in heterozygous mutant brains but levels of *N140fs* transcripts are only approximately 25% of those for wild type transcripts in heterozygous mutant brains (Fig 4). The first round of translation of a transcript is critical for NMD and so inhibition of translation (e.g. with cycloheximide) can increase the stability of transcripts with PTCs (44, 45). Cycloheximide treatment of a group of embryos heterozygous for *N140fs* caused an approximately 5-fold increase in *N140fs* allele-derived transcripts but only an approximately 2-fold increase in wild type transcripts (S4 File) supporting that NMD destabilises *N140fs* transcripts.

**Fig 3.**
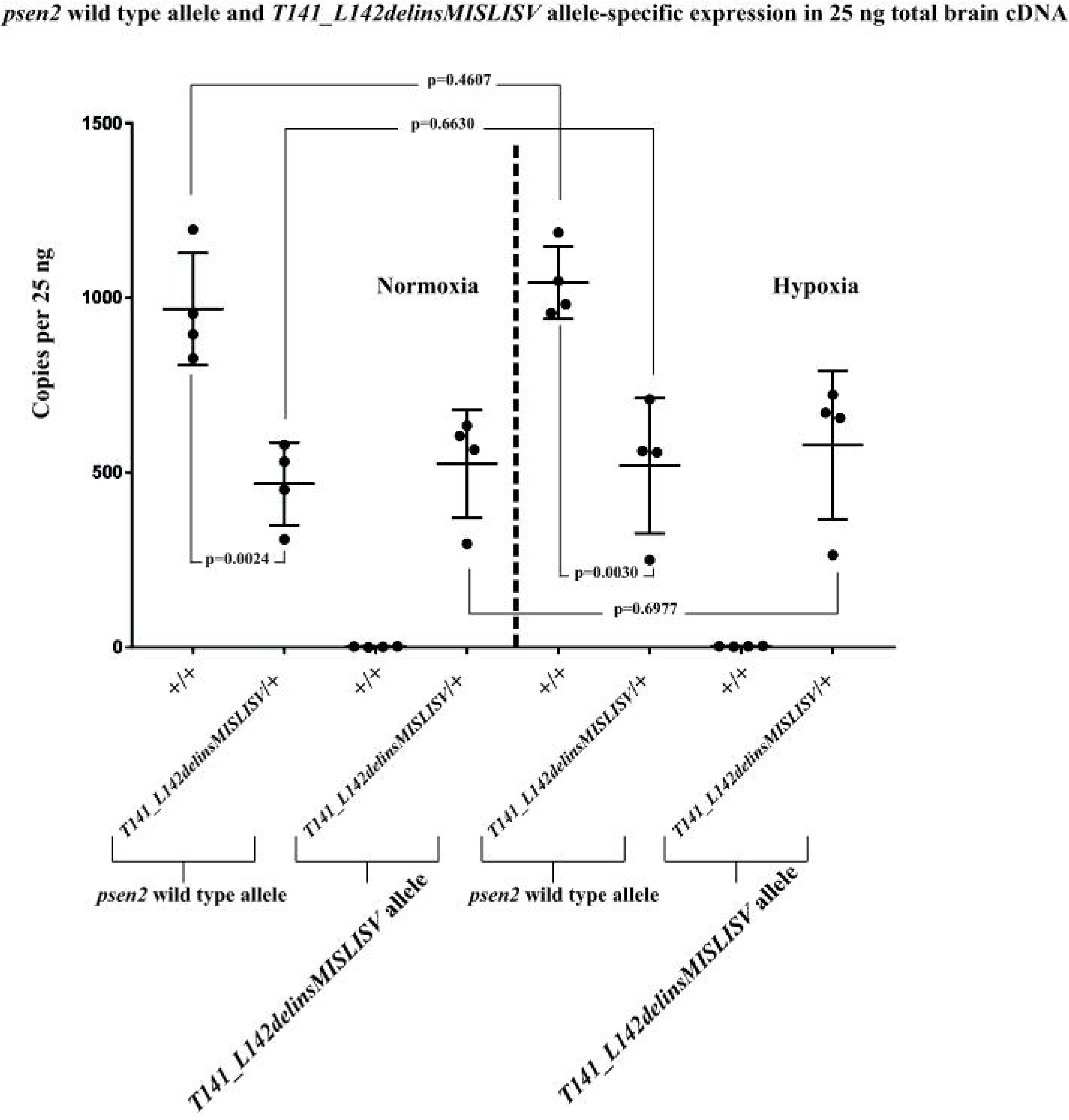
*psen2* wild type and *T141_L142delinsMISLISV* allele-specific expression (as copies per the 25 ng of total brain cDNA in each dqPCR).

**Fig 4.**
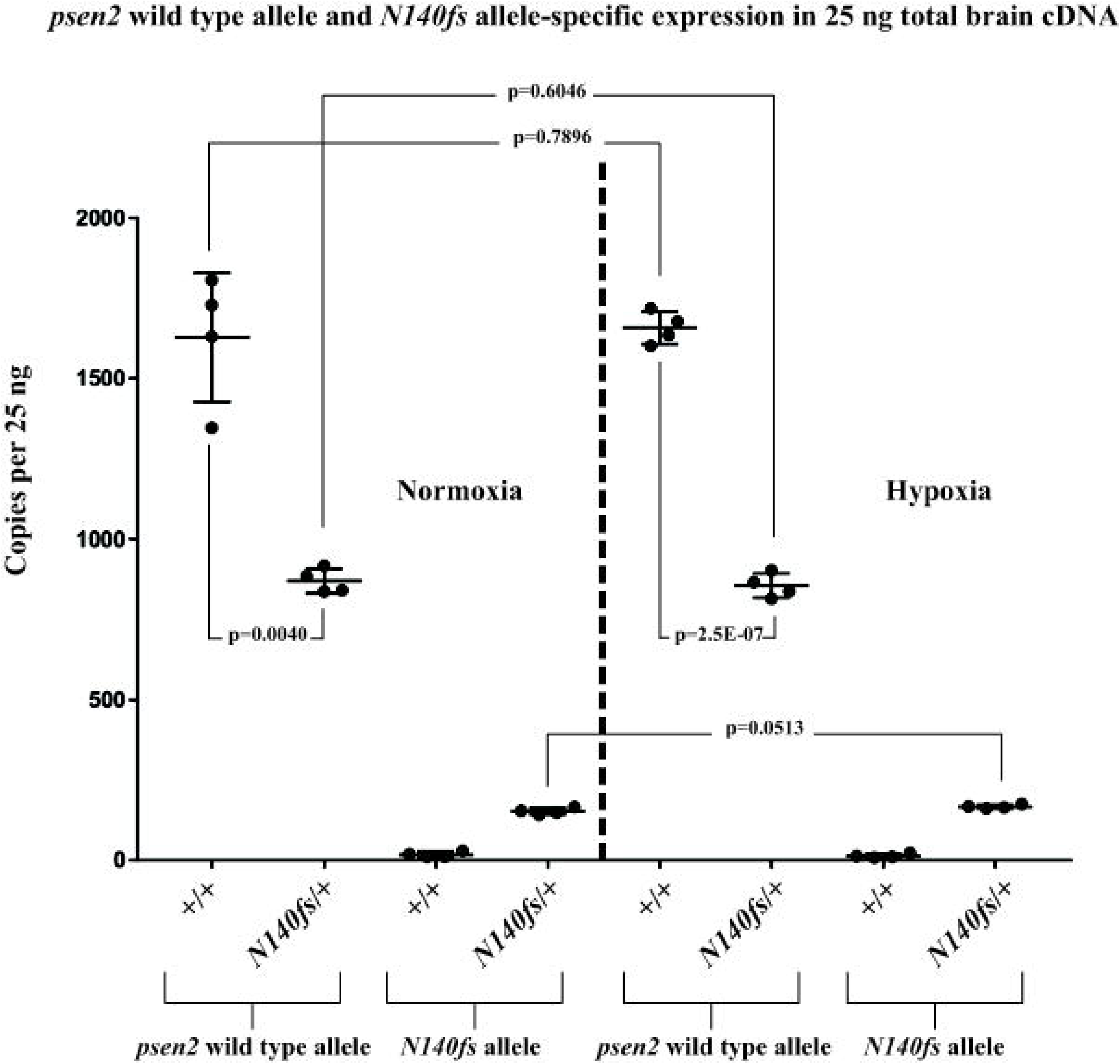
*psen2* wild type allele and *N140fs* allele-specific expression (as copies per the 25 ng of total brain cDNA in each dqPCR). The expression levels of wild type *psen2* alleles in *N140fs*/+ fish (∼860 copies) were significantly (p=0.0024) lower than in their wild type siblings (∼1,600 copies) under normoxia. Under hypoxia, the expression levels of wild type *psen2* alleles in both *N140fs*/+ fish (∼860 copies) and their wild type siblings (∼1,700 copies) were slightly up-regulated, but not with statistical significance compared to these genotypes under normoxia. The expression levels of *N140fs* alleles in *N140fs*/+ fish (∼150 copies under normoxia) were increased (p=0.0513) by acute hypoxia (∼160 copies). Means with SDs are indicated.

### Stability of mutant allele transcripts under normoxia compared to hypoxia

Numerous lines of evidence support that hypoxia is an important factor in the development of AD (reviewed in (46)). This includes that expression of the fAD genes, *PSEN1, PSEN2* and *APP* are upregulated under hypoxia (47-50), phenomena that are conserved in zebrafish (51) despite ∼420 million years of divergent evolution from mammals (52). Also, hypoxia has previously been observed to inhibit NMD (53). Therefore, to observe how hypoxia might affect the levels of transcripts from our mutant alleles we performed dqPCR using total RNA extracted from the brains of 6-month-old zebrafish exposed to normoxia or hypoxia (Figs 3 and 4). This revealed little effect of hypoxia on the levels of transcripts from wild type or *T141_L142delinsMISLISV* alleles (Fig 3) in heterozygous fish brains (that is most likely due to the young age of the fish, see Discussion) and a small, but apparently statistically significant increase in the levels of *N140fs* allele transcripts (Fig 4). However, we cannot distinguish whether this increase is due to induction of transcription, or inhibition of NMD, or both (or other factors that could increase transcript levels).

The expression levels of wild type *psen2* alleles in *T141_L142delinsMISLISV*/+ fish (∼460 copies) were significantly (p=0.0024) lower than in their wild type siblings (∼950 copies) under normoxia. Under hypoxia, the expression levels of wild type *psen2* alleles in both *T141_L142delinsMISLISV*/+ fish (∼1,000 copies) and their wild type siblings (∼510 copies) were up-regulated, but neither of the genotypes showed statistically significant differences compared to their normoxic controls. The expression levels of the *T141_L142delinsMISLISV* alleles in *T141_L142delinsMISLISV*/+ fish (∼520 copies under normoxia) were increased by acute hypoxia (∼580 copies), but without statistical significance. Means with SDs are indicated.

### Pigment phenotypes of mutation-carrying fish

During the process of isolating mutations in *psen2*, we observed that some of the G0 CRISPR Cas9-injected, mosaic, mutation-carrying fish showed unique patches of pigmentation loss in their skin (Fig 5A). (Four of 12 G0 fish injected with the CRISPR Cas9 complex targeting the N140 codon showed this phenotype). None of the F1 progeny of these fish (heterozygous for either of the mutations in *psen2*) showed obvious pigmentation loss. However, when inbreeding F2 heterozygous mutant fish we found that some of the F3 progeny for either the *T141_L142delinsMISLISV* mutation or the *N140fs* mutation showed reduction in surface melanotic pigmentation obvious to the unaided eye by one month of age. Genotyping of these fish using allele-specific PCR on tail biopsies showed them to be homozygous mutants, supporting that the reduced pigmentation phenotypes of the mutations are recessive (see Fig 5B,C). Subsequently, we observed the development of surface pigmentation with age for these fish families and saw that fish heterozygous for either the *T141_L142delinsMISLISV* or *N140fs* mutation appear similar to wild type fish in surface pigmentation but that homozygous *T141_L142delinsMISLISV* fish have much fainter melanotic pigmentation, with many faintly melanotic cells arranged in apparently normal stripes (Fig 5B). In contrast, homozygous *N140fs* fish apparently lack surface melanotic stripes (although a very faint impression of striping is still visible, Fig 5C). Subsequent generation of homozygous lines of fish homozygous for both mutations showed that their reduced melanotic pigmentation phenotypes are consistent and that the mutations do not cause sterility. Since γ-secretase activity is required for melanin formation (20, 25), and *psen2* appears relatively highly expressed in melanophores (33) it is likely that *N140fs* homozygous fish lack melanin due to absence of γ-secretase activity from *psen2* while *T141_L142delinsMISLISV* homozygous fish retain low levels of *psen2*-derived γ-secretase activity.

**Fig 5.**
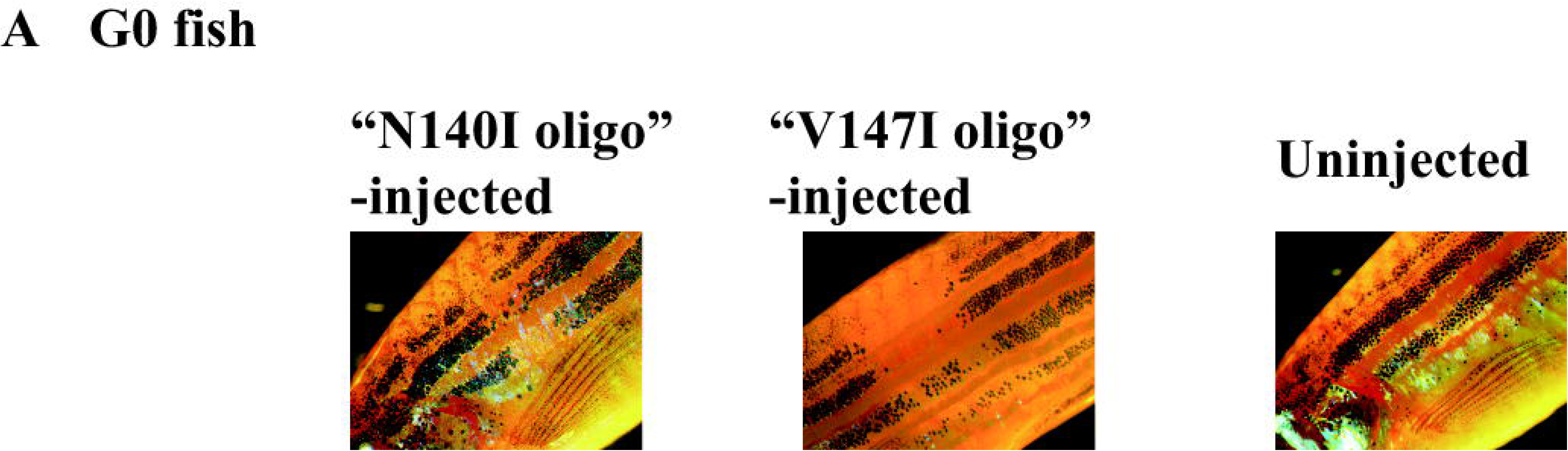
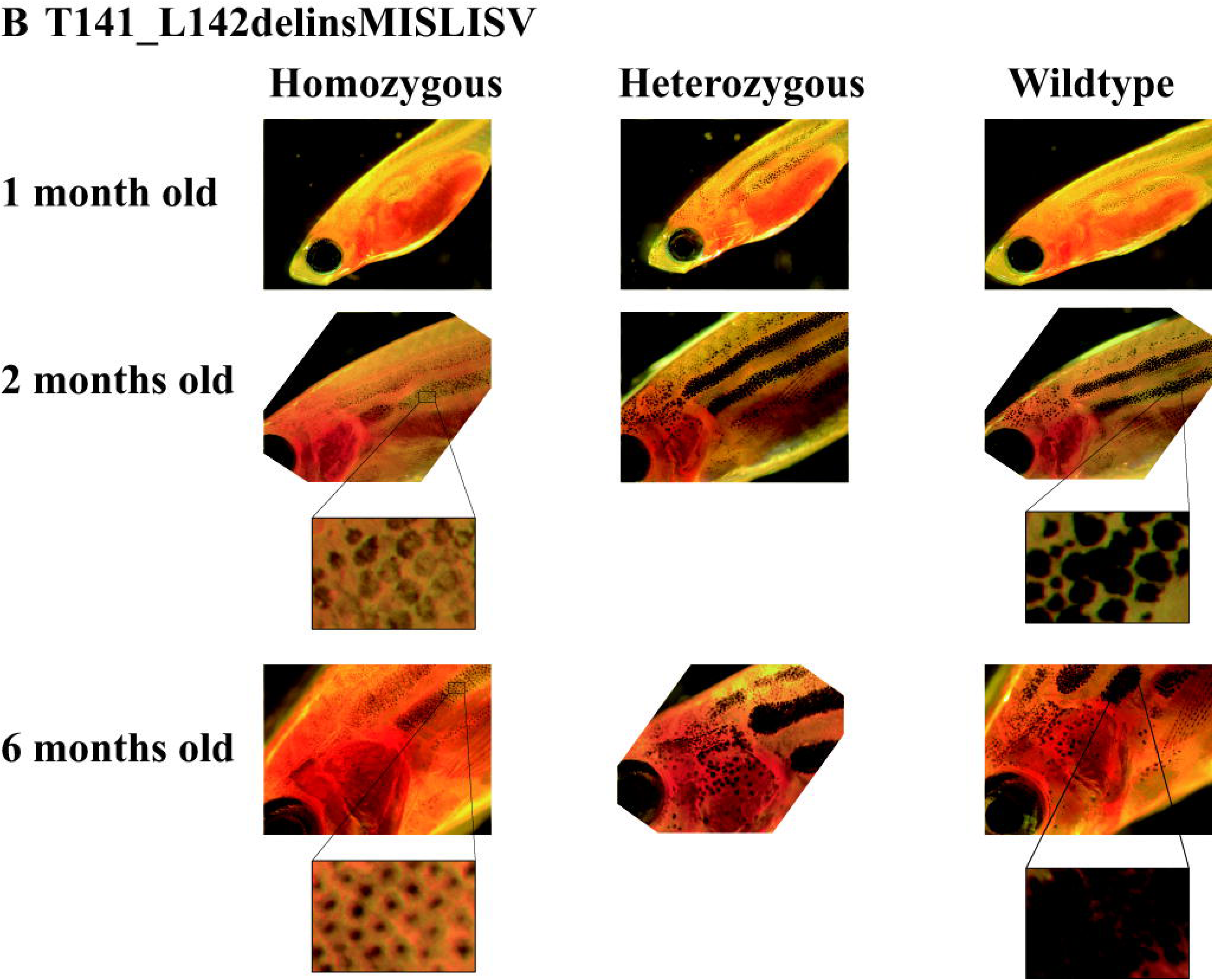
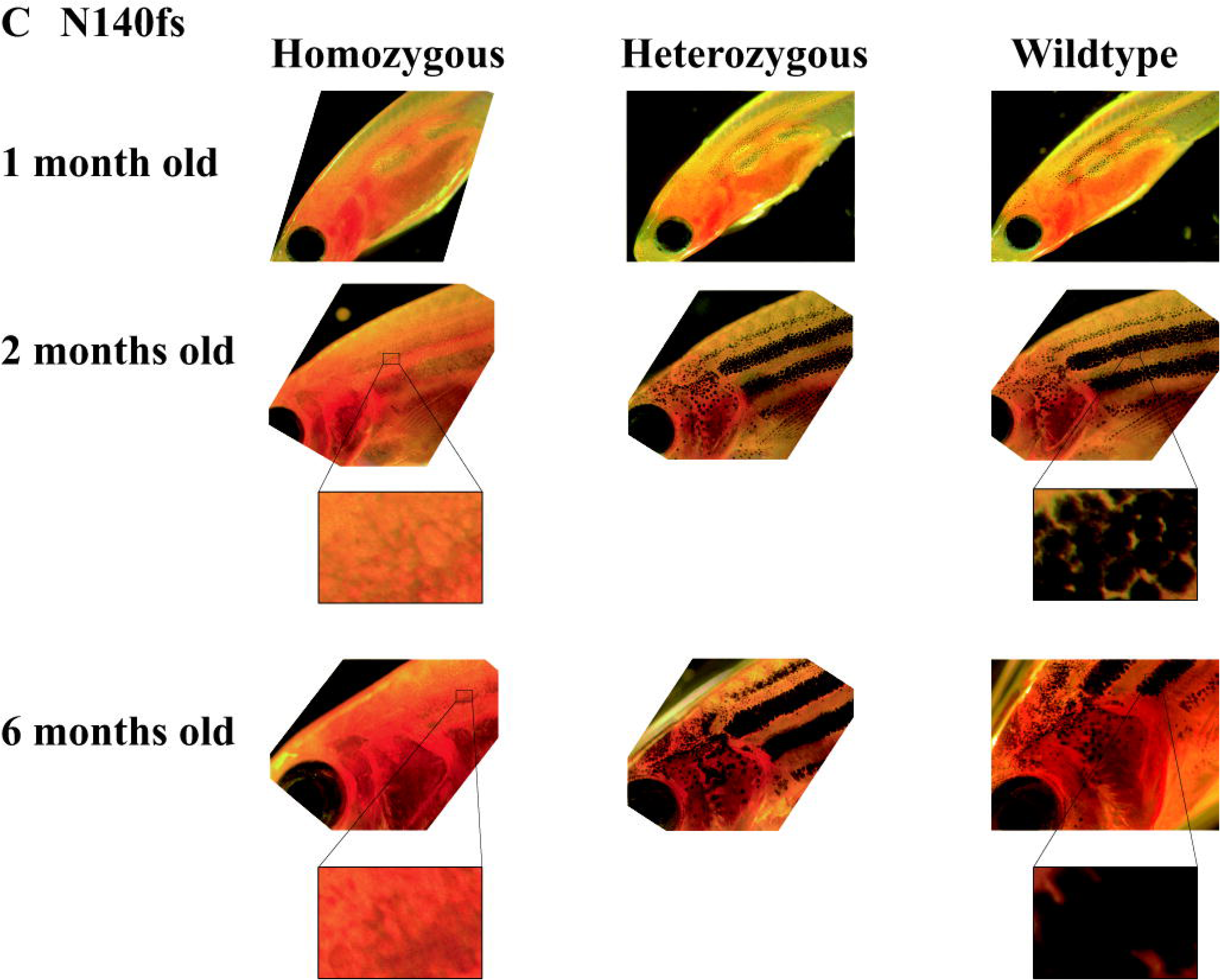
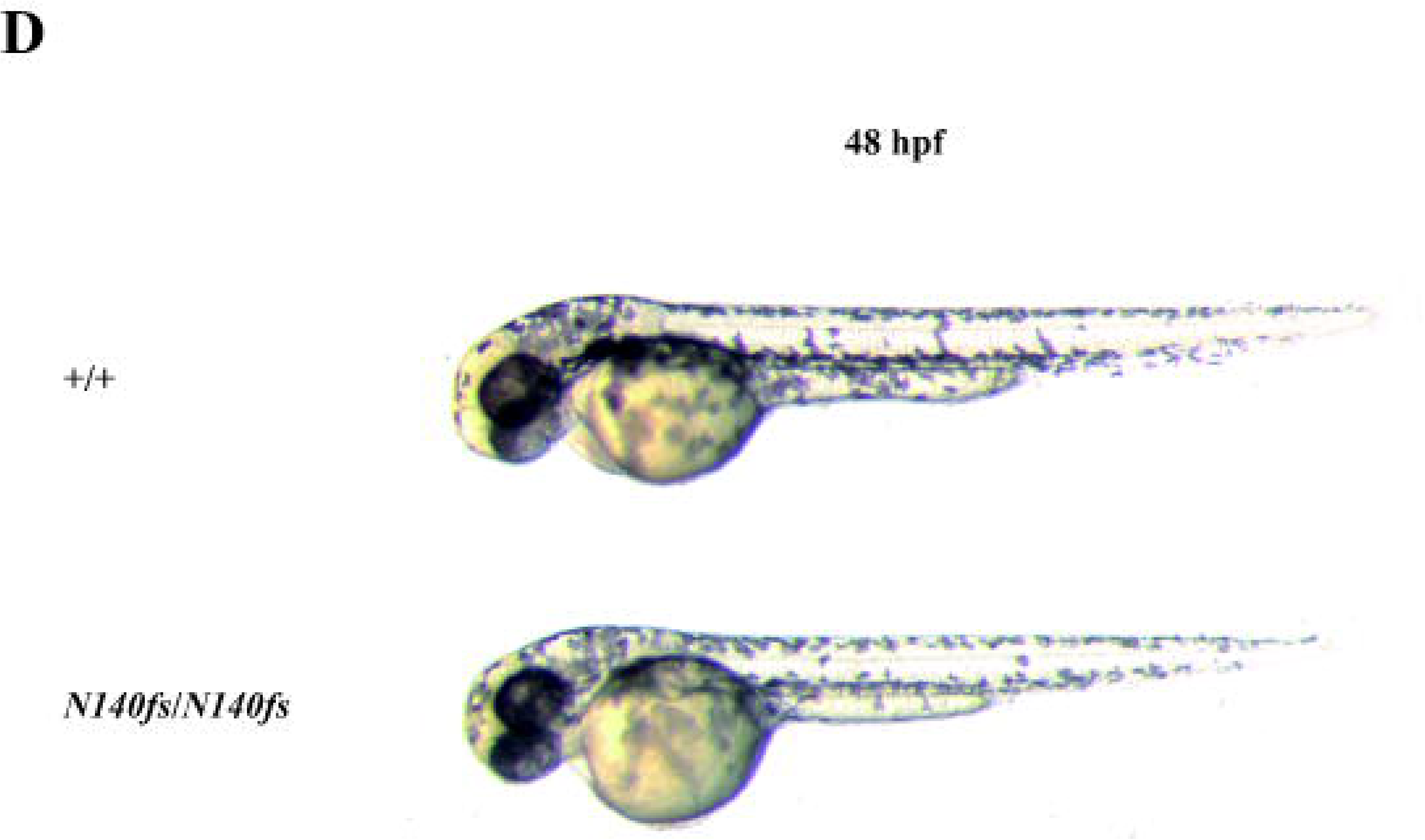
Surface melanotic pigmentation phenotypes. (A) Patches of pigmentation loss in the skin of mosaic mutant G0 fish. (B) *T141_L142delinsMISLISV* mutants and +/+ sibling fish. (C) *N140fs* mutants and +/+ sibling fish. (D) No gross melanotic pigmentation phenotype was observed in *N140fs* homozygous embryos at 50 hpf.

When stripes of melanotic pigmentation were visible in heterozygous or homozygous mutant fish, we did not observe obvious differences in the overall pattern of striping between these and wild type fish (data not shown).

The intracellular distribution of pigment also appeared to change with age in the skin melanophores of *T141_L142delinsMISLISV* homozygous fish. At two months of age pigment appeared evenly distributed in these cells but excluded from their central, presumably nuclear, regions (Fig 5B). However, by six months of age, the pigment appeared concentrated at the centre of cells and was, presumably, perinuclear. The density of pigment formation in heterozygous and wild type fish made it difficult to see whether a similar phenomenon was also occurring in those.

Curiously, the *N140fs* homozygous fish lacking surface melanotic pigmentation retained strong melanotic pigmentation in their retinal pigmented epithelium. This is obvious as the dark eyes of the fish shown in Fig 5C and was confirmed by dissection of these eyes (not shown). Also, all the 48 hpf larval *N140fs* homozygous progeny of homozygous parents showed abundant surface melanophores that cannot be due to maternal inheritance of wild type *psen2* function (Fig 5D). Thus, the dependence of zebrafish adult skin melanotic pigmentation on *psen2* function is both cell type- and age-specific.

## Discussion

Our attempts at generation of point mutations in the zebrafish *psen2* gene by HDR were unsuccessful. However, we did succeed in identifying two mutations (formed by the NHEJ pathway) that may prove useful in analysing the role of the human *PSEN2* gene in familial Alzheimer’s disease; an in-frame mutation, *T141_L142delinsMISLISV*, and a frame-shift mutation, *N140fs*. While the CRISPR Cas9 system can produce off-target effects (54) these are unlikely to have influenced out results since use of this system in zebrafish requires outbreeding of fish that typically segregates away second-site mutations (other than those tightly linked to the target mutation site). Also, the severity of the phenotypic effects observed corresponds to the severity of the effects of the mutations on the structure of the putative encoded proteins. It is also unlikely that two off-target mutation events would both affect pigmentation. Lastly, the effects of mutations in zebrafish *psen2* upon pigmentation are consistent with what is known about the subcellular localization of PSEN2 protein in mammalian systems (10).

The in-frame mutation *T141_L142delinsMISLISV* is an indel mutation altering two codons and inserting an additional 5 codons. Although this mutation changes the length of the protein coding sequence, the predicted protein hydropathicity plot of the putative mutant protein (Fig 2) supports that the mutation does not completely destroy the transmembrane structure of Psen2. Since most of the fAD mutations in human *PSEN2* are in-frame mutations that may change hydropathicity without destroying the overall transmembrane structure of the protein (4), the *T141_L142delinsMISLISV* mutation would appear to be more fAD-like than null.

The frame-shift mutation *N140fs* was caused by a deletion of 7 nucleotides and results in a PTC at the 142^nd^ codon position. This mutation causes truncation of the coding sequence at the upstream end of TMD2 of zebrafish Psen2. The first two TMDs of human PSEN2 are thought to be necessary for ER localisation (11). Since coding sequence truncation occurs at the upstream end of TMD2, if this mutant allele expressed a protein, it would most likely not be able to form TMD structures for ER localization. Neither could it possibly have γ-secretase activity since it lacks the aspartate residues required (41, 42). Moreover, since dqPCR showed that the levels of *N140fs* transcripts are only approximately 25% of those for wild type transcripts in heterozygous mutant brains, *N140fs* expression appears limited by NMD (a fact supported by the ∼5-fold increased N140fs transcript level in the presence of the translation inhibitor, cycloheximide (S4 File). Our previous work has shown that zebrafish *psen2* does not express a truncated isoform equivalent to the PS2V isoform of human *PSEN2* (39) and that a PS2V-like truncation of zebrafish Psen2 does not have PS2V-like activity (55). (Instead a PS2V-like function is expressed from zebrafish *psen1* (55)). Therefore, *N140fs* most likely represents a true null (or severely hypomorphic) allele of zebrafish *psen2*, unlike another frameshift mutation, *S4Ter*, that we recently analysed and that shows grossly normal adult pigmentation (Jiang et al., manuscript submitted).

There is a considerable weight of evidence supporting the importance of hypoxia in the development of AD (reviewed by (56)) and zebrafish represent a very versatile system for investigating the effects of hypoxia (39, 51, 55). In human cells, expression of the fAD genes *APP, PSEN1* and *PSEN2* genes can be upregulated by hypoxia (51) and we previously showed that this phenomenon has been conserved during the nearly half a billion years since the divergence of the zebrafish and human evolutionary lineages (51). In that earlier paper we saw nearly a two-fold increase in zebrafish brain *psen2* mRNA levels under hypoxia compared to normoxia while, in this work, no significant differences were seen (except for *N140fs* allele transcripts where hypoxia may be inhibiting NMD (53)). Upon checking our laboratory records we found that the fish used in the earlier publication were around 12 months old compared to the six months of age in this work. In other, yet unpublished work we have observed that differences in adult age make very significant differences to brain transcriptional responses to hypoxia with young adult fish showing the mildest responses (Newman et al. unpublished results).

In previous research we showed that blockage of *psen2* function using morpholino antisense nucleotides injected into zebrafish zygotes increases the number of DoLA neurons at 24 hpf (57). Despite the evidence that the *N140fs* mutation is null, we did not see increased DoLA neuron numbers in *N140fs* homozygous embryos at 24 hpf (See S3 Files for experiment description and data). The observation of differing developmental phenotypes from decreased gene function due to mutation or morpholino injection is a common occurrence (58). It is thought to be due to the phenomenon of “genetic compensation” whereby only decreased gene function through mutation, (and not by morpholino injection), causes compensatory upregulation of other genes with similar activities (59). It is likely that genetic compensation is causing the lack of response of DoLA neuron number to the *N140fs* mutation. An alternative explanation would be a maternal contribution of wild type *psen2* activity from the heterozygous *N140fs* mother of the embryos examined. Further experimentation such as blockage of *psen1* translation by morpholino injection into *N140fs* homozygous embryos or analysis of DoLA numbers in *N140fs* homozygous embryos from homozygous parents might resolve this question.

The visually striking surface pigmentation pattern of zebrafish and the genetic utility of this organism has made it a focus for research on the genetic control of pigment formation (60), pigment cell differentiation (61), and surface pigmentation pattern formation (62). Skin pigmentation pattern is severely affected in adult fish homozygous for the mutation *T141_L142delinsMISLISV*. These fish show surface melanotic stripes that appear approximately the same width as in wild type fish but are much fainter. Closer examination of these stripes at 6 months of age reveals cells with vestigial, and likely perinuclear, pigment. The number of cells is not obviously affected, only the pigmentation they show. Thus, loss of *psen2* function does not appear to affect melanophore viability (although, in an animal as highly regenerative as the zebrafish, further tests would be required to conclude this with certainty). By extrapolation it appears likely that *N140fs* homozygous adult fish still possess skin melanophores but that these lack melanin. The retention of some adult skin melanin formation in *T141_L142delinsMISLISV* homozygotes but not *N140fs* homozygotes, and the roles played in melanosome formation and function particularly by PSEN2-derived γ-secretase activity (10, 28), support that *T141_L142delinsMISLISV* mutant Psen2 protein molecules retain some level of γ-secretase activity. (Indeed, Sannerud et al (10) observed that loss of *PSEN2* activity in a human melanoma cell line, MNT1, greatly reduced γ-secretase cleavage of tyrosinase-related protein (TRP1) and premelanosome protein (PMEL) that are important for melanosome function.) This supports that the *T141_L142delinsMISLISV* mutation of zebrafish Psen2 does not seriously disrupt the protein’s overall pattern of folding for membrane insertion, but does distort its conformation sufficiently to reduce γ-secretase activity. Partial loss of γ-secretase activity is a commonly observed characteristic of fAD-like mutations in *PRESENILIN* genes. For example, mouse skin completely lacking expression of wild type *Psen1* and *Psen2* genes but with a single knock-in M146V fAD-like allele of *Psen1* show lighter skin and coat colour than similar mice possessing a single wild type allele of *Psen1* (20). These data, and the fact that the *T141_L142delinsMISLISV* mutation obeys the “fAD mutation reading frame preservation rule” (7), support that this mutation should be investigated for its utility in zebrafish-based fAD research.

Intriguingly, only the melanotic pigmentation of adult zebrafish skin is dependent on *psen2* function while larvae and cells of the retinal pigmented epithelium do not show this dependency. In mammalian systems, most melanin synthesis in the retinal pigmented epithelium occurs during embryogenesis (63). However, maternal inheritance of wild type *psen2* mRNA acting during zebrafish embryo formation cannot explain the pigmented melanophores of larvae or the pigmentation in adult retinas of *N140fs* homozygotes since this pigmentation is observed in the progeny of homozygous mutant parents. The pigmentation likely indicates that the Psen1 protein (or, possibly, another protein with γ-secretase-like activity (7)) contributes to normal melanosome formation in the melanophores of embryos/larvae and in the retinal pigmented epithelium of zebrafish. That different PRESENILIN proteins might contribute differentially to melanosome formation in different cells or in the same cell type at different ages is a level of developmental complexity that has not previously been appreciated. Alternatively, the skin melanophores of adult fish might, for some unknown reason, be incapable of genetic compensation (e.g. upregulation of *psen1* activity when *psen2* activity is lost through mutation). The possibility of cell type-specificity of genetic compensation has also not previously been considered. The lack of an obvious larval pigmentation phenotype explains why *psen2* was not identified by the large mutation screens for developmental phenotypes conducted by the laboratories of Christiane Nüsslein-Volhard (64) and Wolfgang Driever (65) and published in 1996.

In conclusion, we have generated in zebrafish an EOfAD-like mutation, *T141_L142delinsMISLISV*, and an apparent null, loss-of-function mutation *N140fs*. Since none of the over 200 human fAD mutations in *PSEN1* and *PSEN2* are obviously null alleles, these two zebrafish mutations may prove useful for defining the brain gene regulatory and other molecular changes that are particular to fAD mutations in the *PRESENILIN* genes. Our future work will use these and other zebrafish mutation models to dissect how fAD-like mutations contribute to Alzheimer’s disease. Also, Higdon et al (66) showed that it is possible to use cell-sorting techniques on disassociated zebrafish embryos to isolate relatively pure populations of their different pigment cells types. These were subsequently characterised transcriptomically. Extension of these technologies to larval, retinal, and adult tissues would allow more detailed analysis of the differences between the melanotic cells of these stages and tissues to determine why they are differentially dependent on *psen2* activity for melanosome formation and function.

## Acknowledgments

The authors wish to thank Seyyed Hani Moussavi Nik for kind assistance in adjusting the conditions for the dqPCR.

## Supporting information

**Fig. S1.**
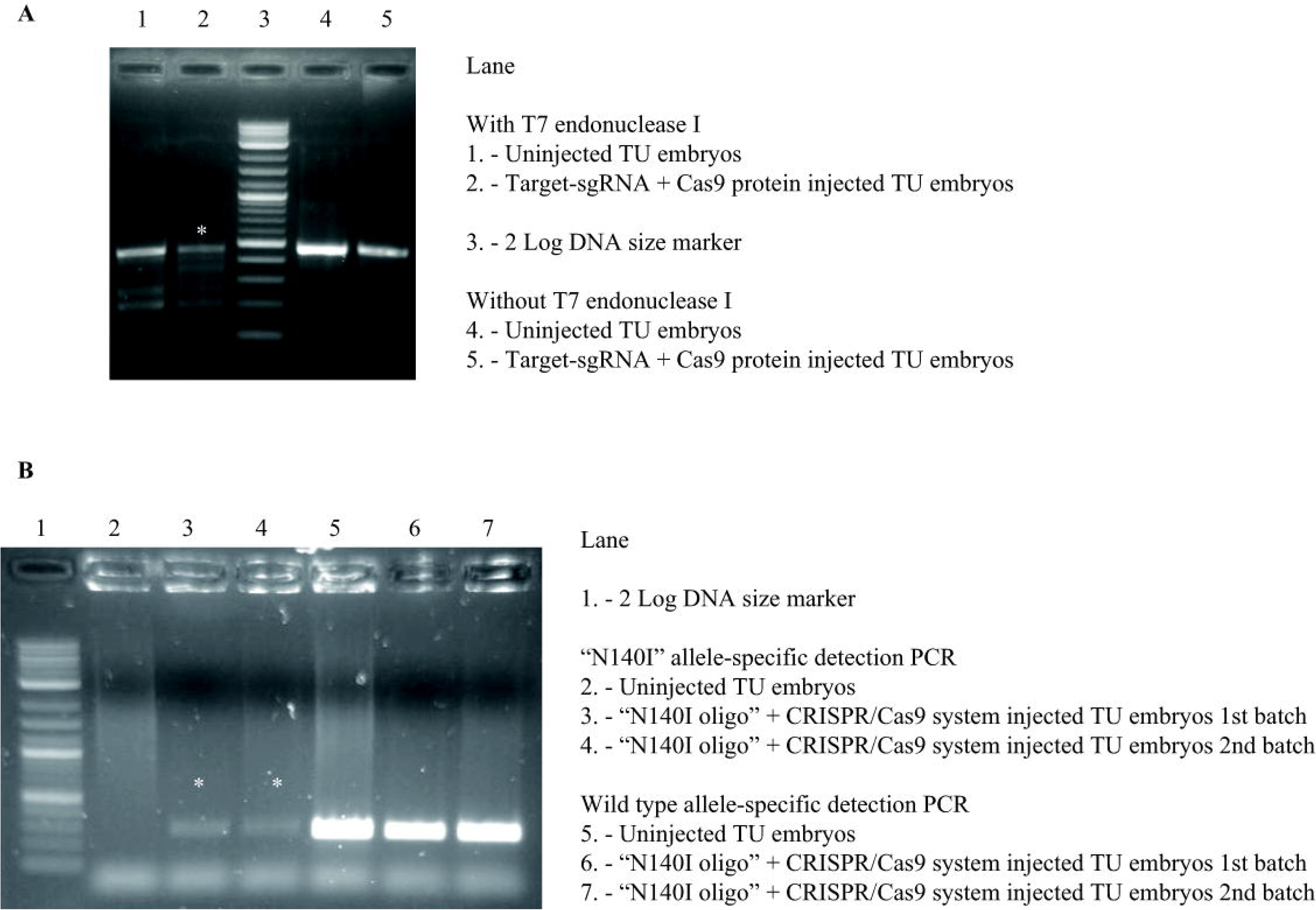
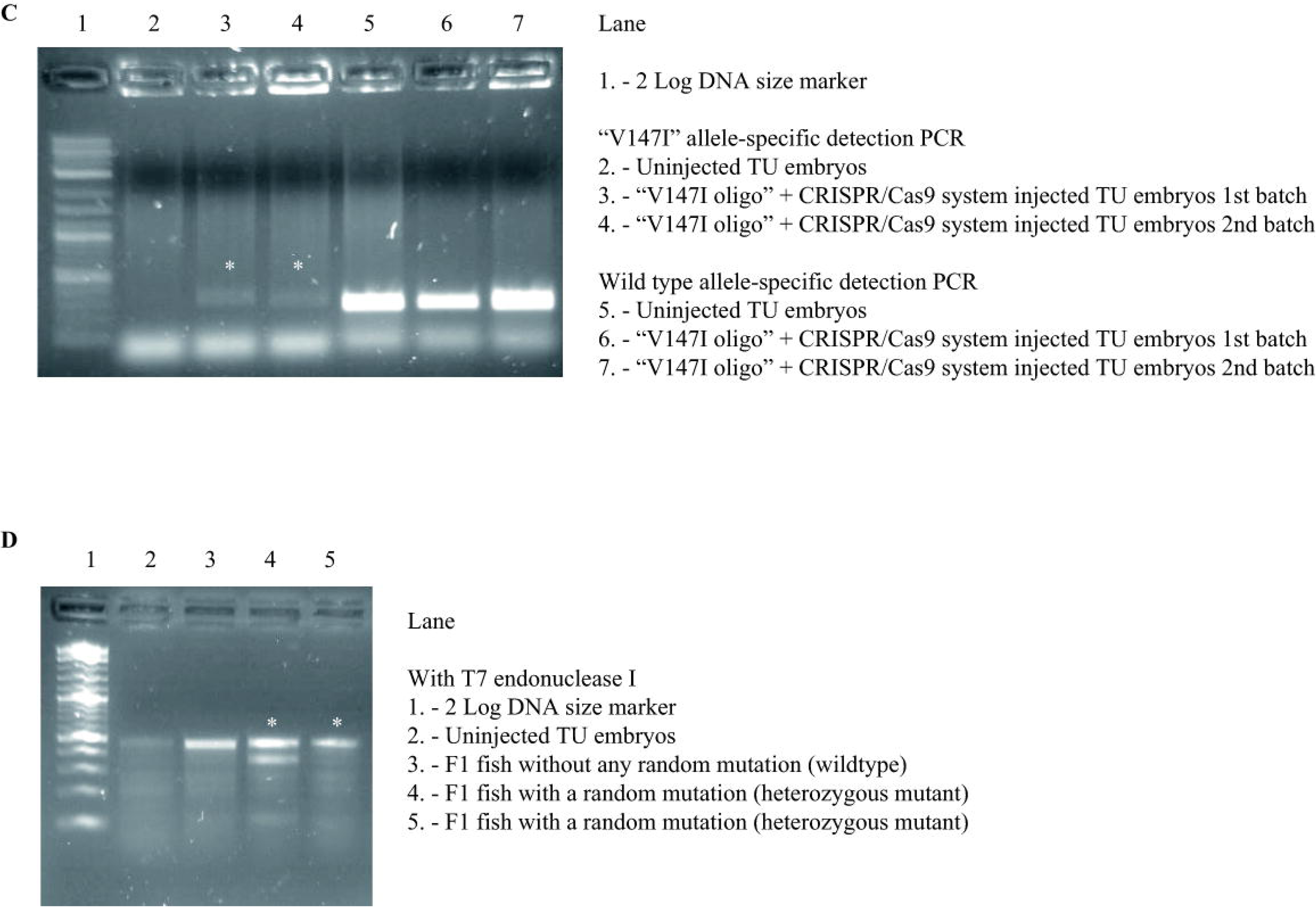
T7 endonuclease assays and mutation-specific PCRs for embryos at 24 hpf. (A) T7 endonuclease I assay for testing the cleavage activity of the CRISPR/Cas9 system.(B) “*N140I*” allele-detection PCR for testing of CRISPR/Cas9 plus “N140I oligo” co-injected TU embryos. 10 embryos from each injection batch were pooled for these tests. Both batches of the injected TU embryos showed positive signals in the “N140I” allele-detection PCR. Therefore, some of these “N140I oligo” injected TU embryos may have carried the “N140I” allele in the genomes of some cells.(C) “*V147I*” allele-detection PCR for testing the CRISPR/Cas9 plus “V147I oligo” injected TU embryos. 10 embryos from each batch were pooled for these tests. Both batches of the injected TU embryos showed positive signals from the “*V147I*” allele-detection PCR. Therefore, some of these “V147I oligo” injected TU embryos may have carried the “*V147I*” allele in the genomes of some cells.(D) T7 endonuclease I assay for detecting random mutations at the CRISPR/Cas9 target site in the F1 progeny. Tail-clip biopsies from 46 of the F1 progeny from the CRISPR/Cas9 plus “V147I oligo” injected mosaic G0 fish were tested using the T7 endonuclease I assay to screen for the presence of cells with mutations at the target site. Only 5 fish showed cleavage patterns indicating the presence of mutations.

**Fig. S2.**
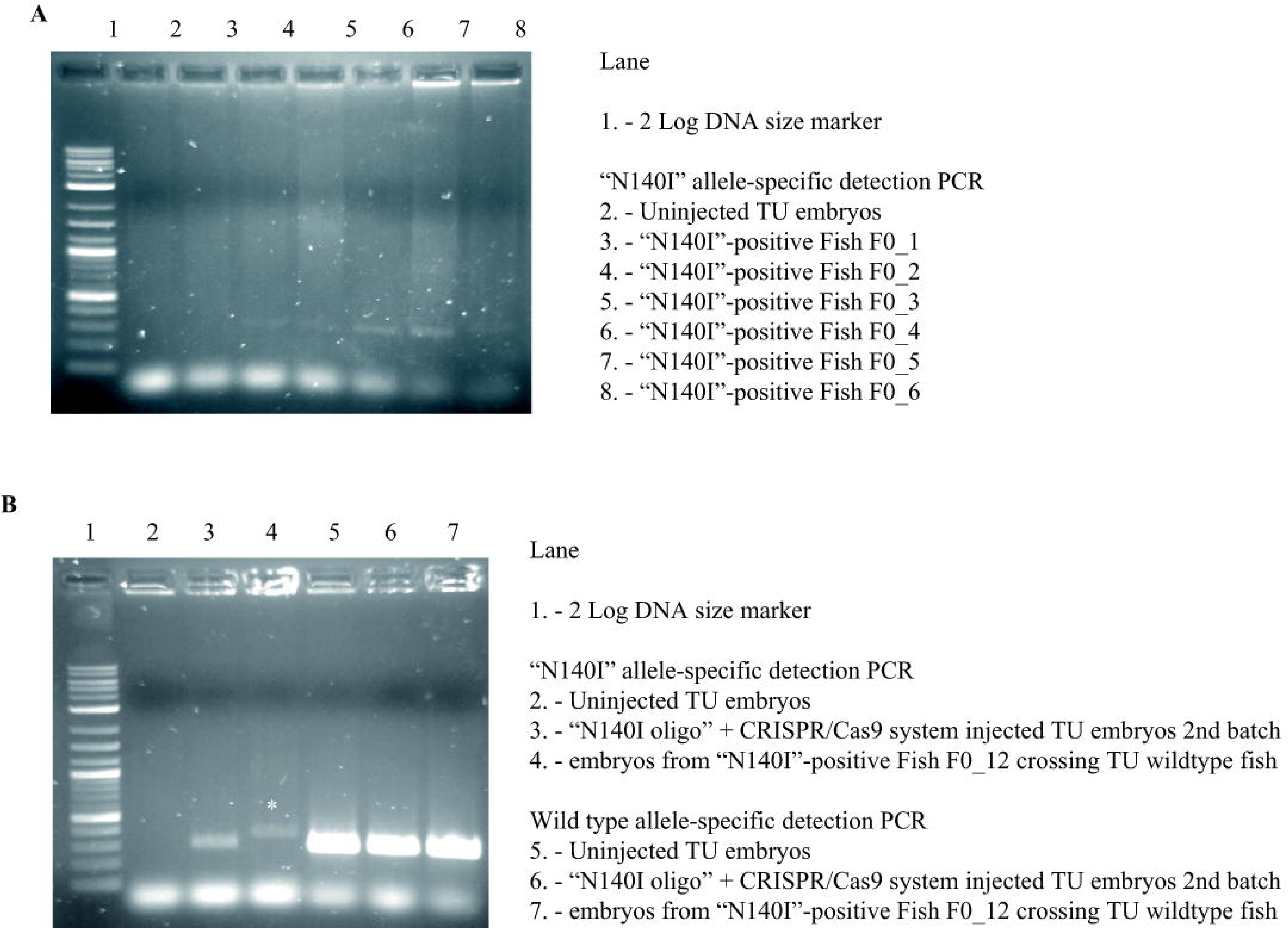
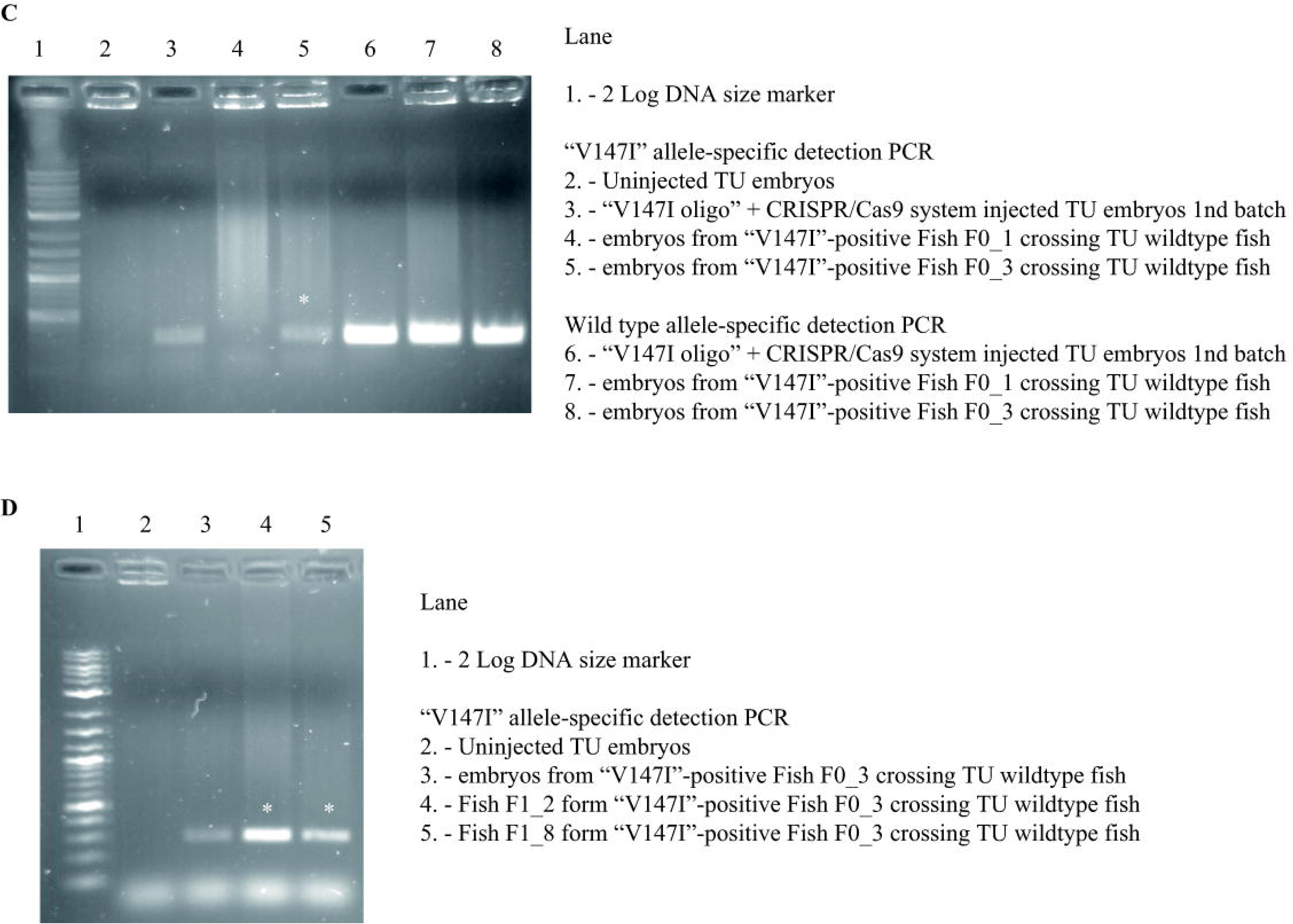
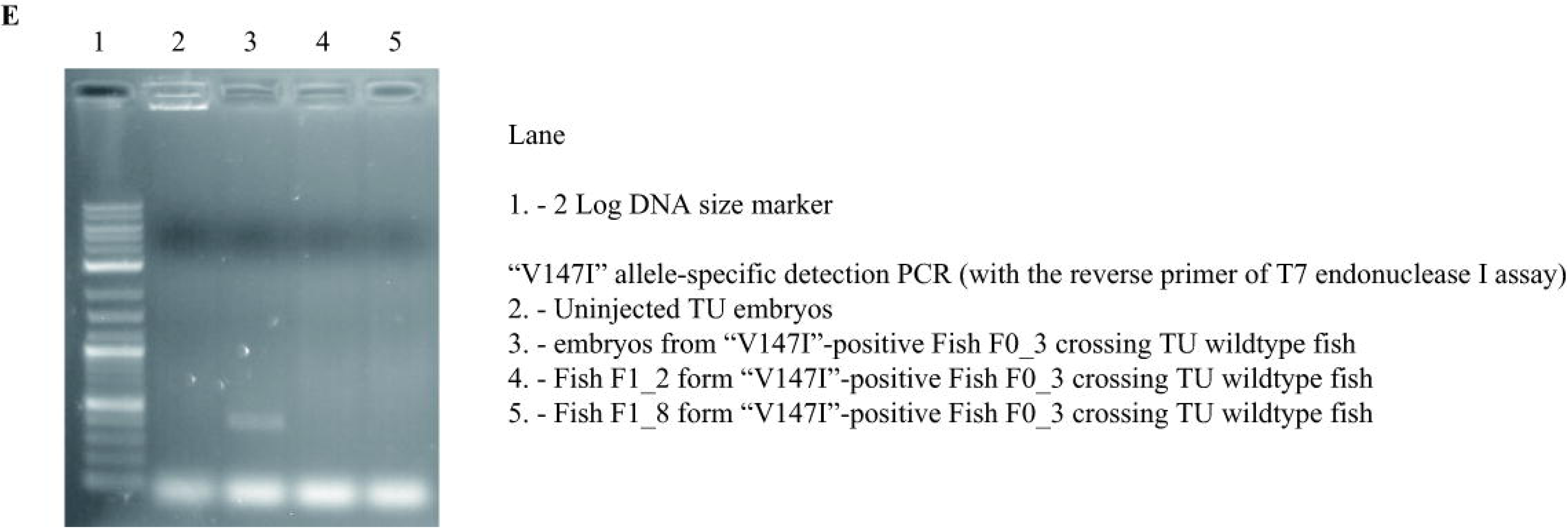
Mutation-specific PCR tests of G0 and F1 fish. (A) “*N140I*” allele-specific detection PCRs on tail-clip biopsies from G0 fish. Twelve G0 fish (120 in total) showed positive signals in the “N140I” allele-specific detection PCR.(B) “*N140I*” allele-specific detection PCRs from F1 embryos of the G0 mosaic fish showing “*N140I*” allele-positive signals. 10 F1 embryos at 24 hpf from each “*N140I*” allele-carrying G0 fish were pooled for testing. The F1 progeny from one “*N140I*” allele-carrying G0 fish showed a signal at ∼400 bp, which may result from imperfect incorporation of the “N140I oligo” sequence into the target site of the CRISPR/Cas9 system.(C) “*V147I*” allele-specific PCRs from F1 embryos of the G0 mosaic fish showing “*V147I*” allele-positive signals. 10 F1 embryos at 24 hpf from each “*V147I*” allele carrying G0 fish were pooled for testing. The F1 progeny from one of the “*V147I*” allele-carrying G0 fish showed the same positive signal as the injected G0 embryos.(D) “*V147I*” allele-specific detection PCR from tail-clip biopsies of F1 fish. Two out of twelve tested F1 fish (raised from the positive batch of embryos observed above in showed positive signals, indicating they might carry the desired “*V147I*” allele.(E) “*V147I*” allele-specific detection PCR using the same forward primer as in (E) but a different reverse primer binding farther downstream in *psen2* DNA. While the pooled F1 embryos still gave a positive signal, the two F1 fish no longer showed a positive signal using this PCR, revealing that the previously seen positive signals (in were artefacts.

**Fig. S3.**
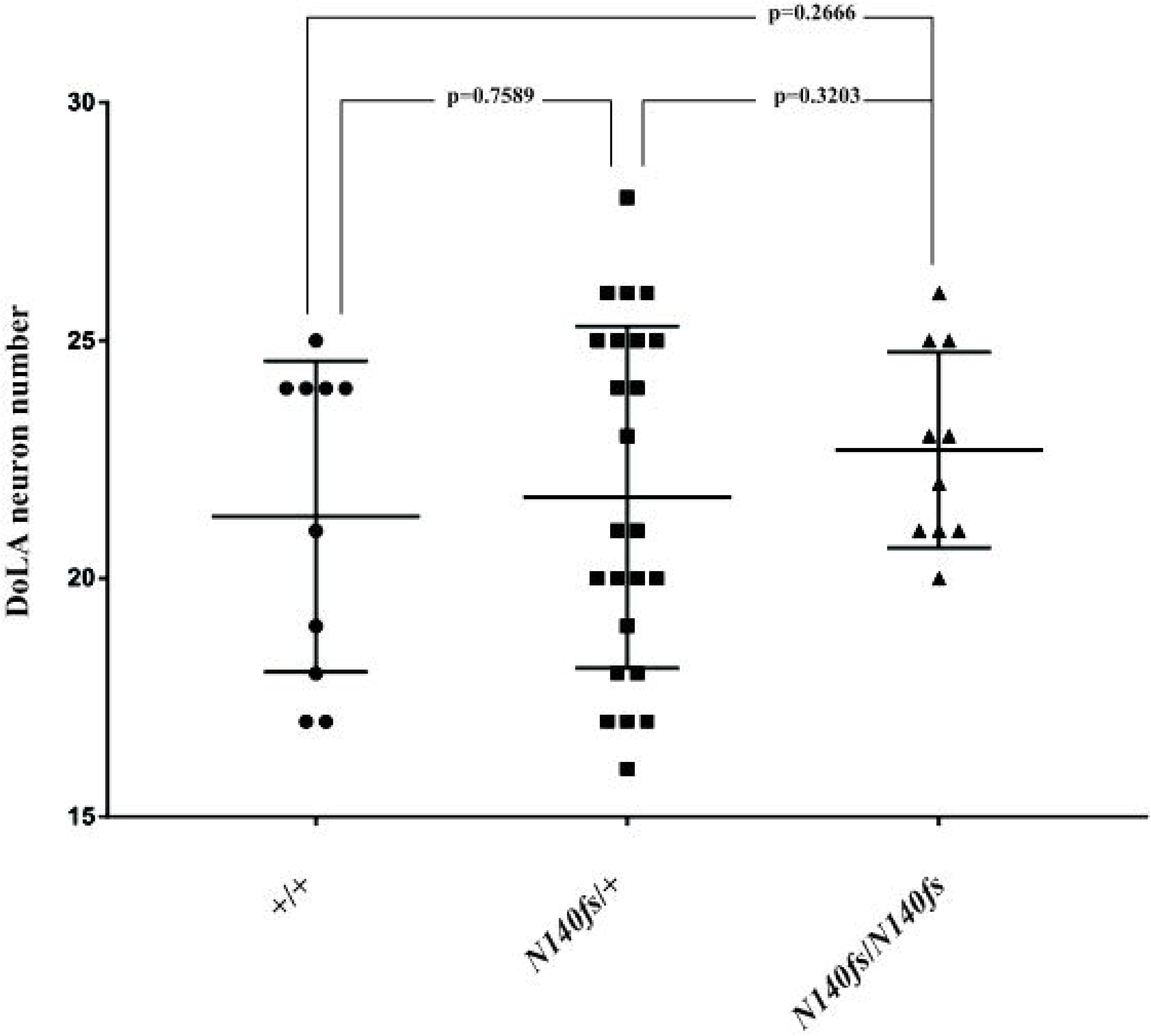
DoLA neuron numbers. DoLA neuron numbers in wild type and *N140fs* mutant embryos as revealed by *in situ* hybridisation against *tbx16* transcripts.

**Fig. S4.**
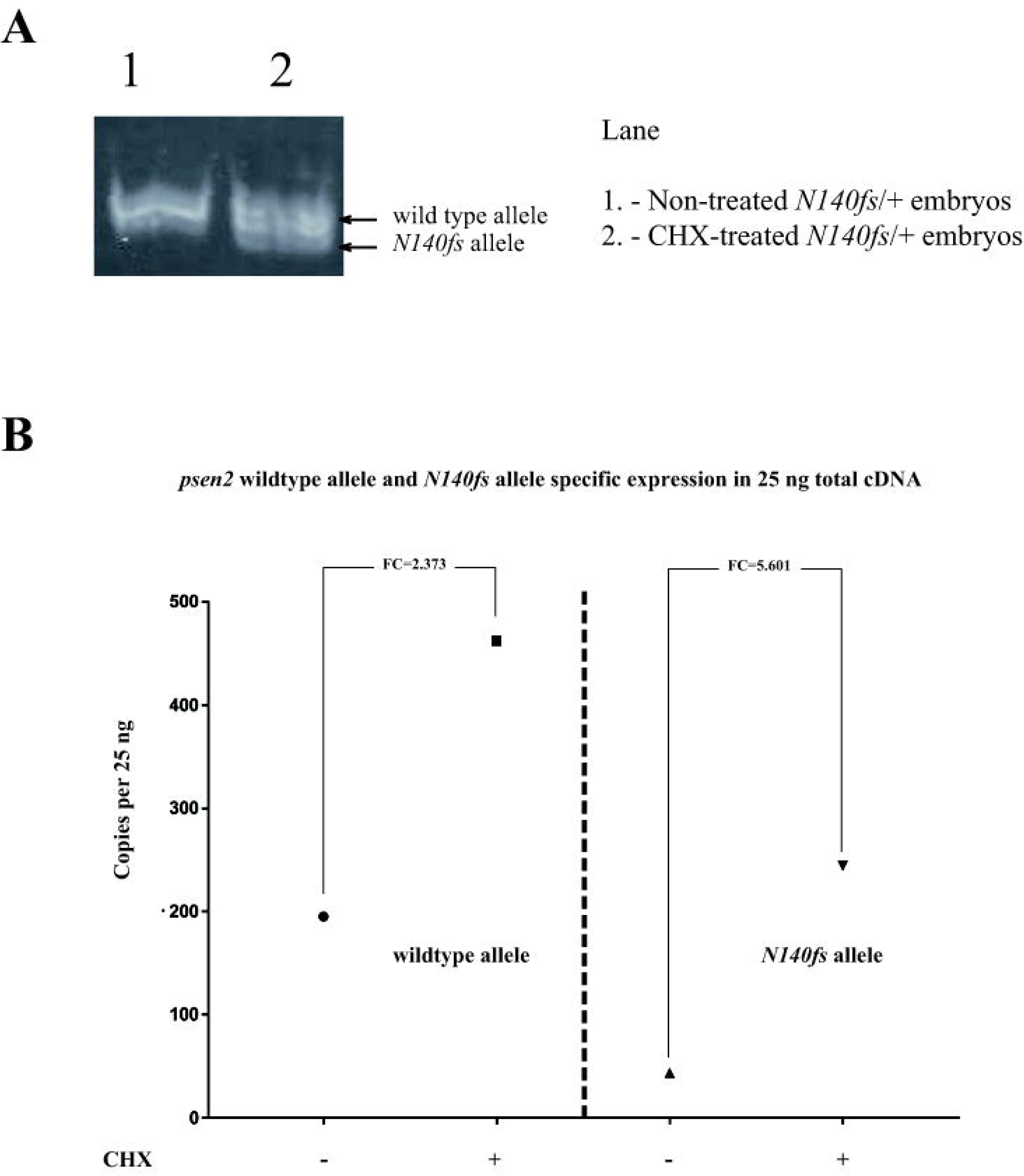
dqPCRs detecting wild type and mutant alleles in *N140fs*/+ embryos at 50 hpf after two hours of cycloheximide treatment relative to untreated embryos. A) In a 20% polyacrylamide gel, amplification of cDNA fragments spanning the mutation (7 nucleotides shorter than wild type) was only observed in the CHX-treated group, while only one higher molecular weight band (from the wild type allele) was observed in the non-treated group. This supports that NMD is destabilising the mutant transcript in heterozygous embryos. In dqPCR, both the wild type *psen2* allele and the *N140fs* allele were observed to be upregulated after the (B) CHX-treatment. The fold change (FC) of the upregulation of the *N140fs* allele transcripts (FC=5.601) was significantly higher than that for the wild type *psen2* allele transcripts (FC=2.373).

**S1 Table. Allele-specific transcript quantification in six month old *T141_L142delinsMISLISV*/+ and wild type sibling brains.** Copies per 25ng of total brain cDNA (assuming complete reverse transcription of total brain RNA).

**S2 Table. Allele-specific transcript quantification in six month old *N140fs*/+ fish and wild type sibling brains.** Copies per 25ng of total brain cDNA (assuming complete reverse transcription of total brain RNA).

**S3 Table. *In situ* hybridization against *tbx16* transcripts in DoLA neurons.**

**S4 Table. Allele-specific expression analysis on the *N140fs*/+ embryos (non-treated and CHX–treated) at 50 hpf in 25ng of total embryo cDNA.** Copies per 25ng (assuming complete reverse transcription of total RNA).

**S1 File. Mutation screening and breeding.**

**S2 File. dqPCR results for allele-specific transcript quantification in six month old brains.**

**S3 File. *In situ* transcript hybridization analysis of DoLA neuron number.**

**S4 File. Cycloheximide treatment of *N140fs*/+ embryos at 48-50 hpf.**

## Abbreviations

AD: Alzheimer’s Disease
CRISPR: clustered regularly interspaced short palindromic repeats
DSB: double-strand break
HDR: homology-directed repair
NHEJ: nonhomologous end joining
NMD: nonsense-mediated decay
PSEN: PRESENILIN
PTC: premature translation-termination codon
TMD: transmembrane domain

